# Constitutive production of flagellar proteins is required for proper flagellation in *Shewanella putrefaciens*

**DOI:** 10.1101/2022.07.21.500047

**Authors:** Meike Schwan, Ariane Khaledi, Sven Willger, Kai Papenfort, Timo Glatter, Susanne Häußler, Kai M. Thormann

## Abstract

Flagella are multiprotein complexes whose assembly and positioning requires complex spatiotemporal control. Flagellar assembly is thought to be controlled by several transcriptional tiers, which is mediated through various master regulators. Here, we revisited the regulation of flagellar genes in polarly flagellated gammaproteobacteria by the regulators FlrA, RpoN (σ^54^) and FliA (σ^28^) in *Shewanella putrefaciens* CN-32 at the transcript and protein level. As expected, strict control at both levels occurred for for highly abundant flagellar proteins, including the building blocks for the outer rings, rod, hook and filament. In contrast, a number of regulatory and structural proteins were always present also in the absence of the main regulators. Initiation of flagella assembly and motor activation likely relies on the abundance control of only few structural key components required for formation of the MS- and C-ring and the flagellar type III secrection system. We identified σ^70^-dependent promoters driving constitutive expression of some flagellar genes including the regulators of flagellar number and positioning, FlhF and FlhG. Reduction of the constitutive expression levels resulted in emergence of hyperflagellation. Thus, basal expression and presence of flagellar proteins is required for proper flagellation, which adds a deeper layer to the regulation of flagellar synthesis and assembly.

**Significance:** The tier-based transcriptional regulation underlying bacterial flagella synthesis is – with certain variations – well-established in various species. Here we show that initiation and proceeding of flagellar synthesis can be simply based on the control of some key components and highly abundant building blocks. We further identified a ‚tier zero’, a set of constitutively produced flagellar regulators and building blocks, which is required, for example, to maintain the flagellar counter. We expect this not only to apply to our model species *Shewanella*, but also to other flagella regulation systems in bacteria.

## Introduction

Many bacterial species are able to synthesize flagella, long proteinaceous helical fibers that extend from the cell’s surface. Rotation of the flagellar helix, mediated by a membrane-embedded motor, enables efficient active movement of the cell through liquid environments (swimming) or across surfaces (swarming) [1–3]. In addition, the flagellum can serve as an adhesive or pathogenicity factor or as a sensor the bacteria use to determine environmental conditions such as viscosity or surface wetness [4,5].

The flagellum is an intricate nanomachine, which - in its general design - is well-conserved between different bacterial species. It consists of the helical filament, the cell envelope-embedded basal body housing the rotary motor and a type III export apparatus, and a universal joint structure, the hook, which connects motor and filament (**Fig. 1A; Supplementary Fig. 1A**) [6,7]. The whole structure is formed from about 20 different protein building blocks with different stoichiometries ranging from less than 10 copies, e.g. for parts of the flagellar type III export system (fT3SS), to tens of thousands of the main filament building block, the flagellin [8,9]. The flagellar motor is rotated at the expense of transmembrane ion gradients, most motors either use H^+^ or Na^+^ as coupling ions [10]. As synthesis, assembly and operation requires a considerable amount of energy and cellular resources, formation and function of flagella are tightly regulated.

**Figure 1:**
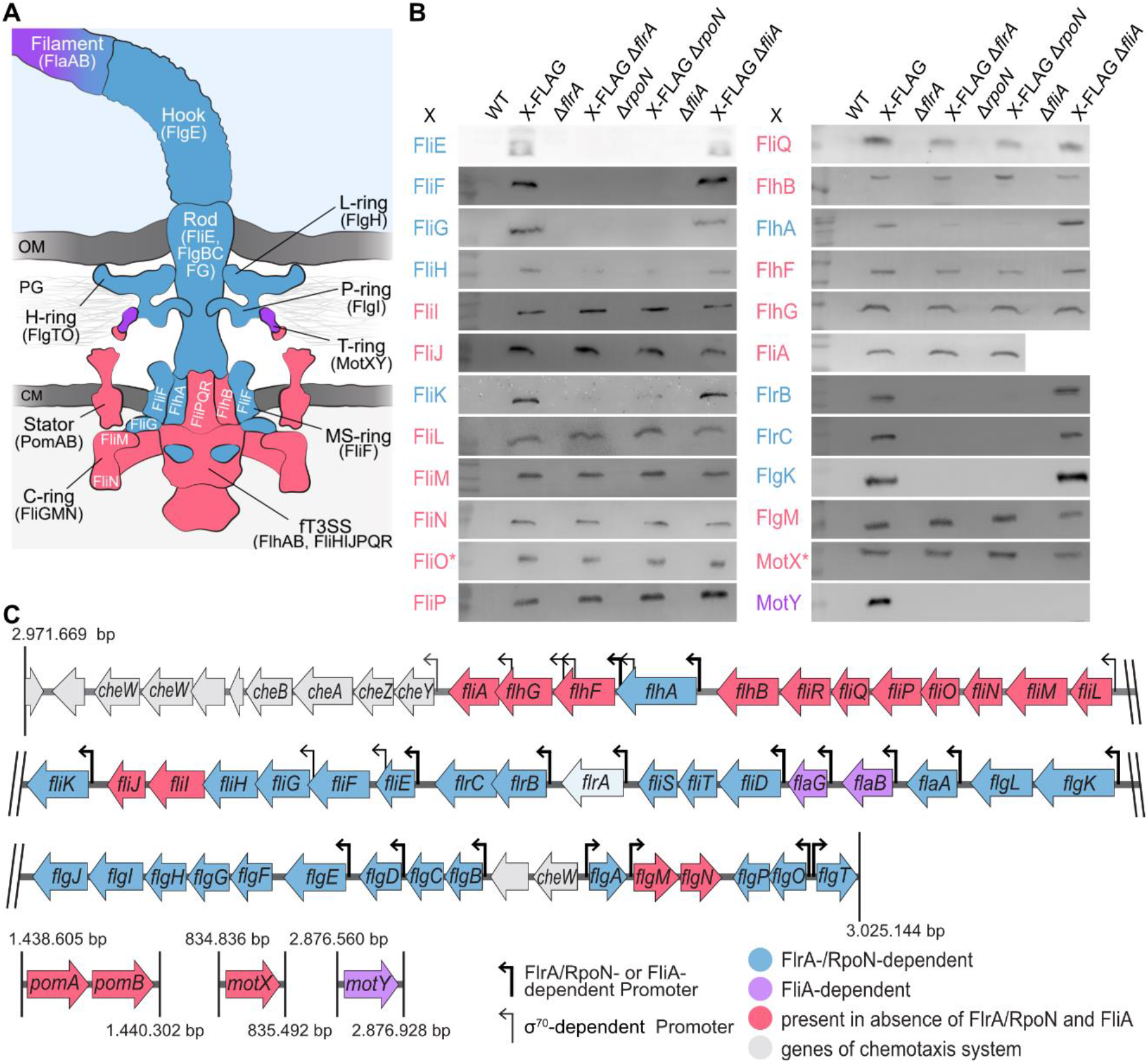
Presence of flagellar components in the absence of flagellar regulators. Flagellar components marked in red are also present in the deletion mutants of the regulators *flrA, rpoN* and *fliA*. Blue-labeled components are FlrA-as well as RpoN-dependent, and purple-labeled components are FliA-dependent. **(A)** Schematic representation of the flagellum (modified from [64]). OM, outer membrane; PG, peptidoglycan, CM, cell membrane. **(B)** Western blot analyses of some flagellar components in deletion strains of the flagellar master regulator *flrA* or the sigma factors *rpoN* and *fliA*, respectively. The corresponding proteins were chromosomally fused with a 3xFLAG, sfGFP (FliO), or mCherry (MotX). The complete blots as well as Coomassie-stained SDS polyacrylamide gels serving as loading controls are shown in **Supplementary Figures 4 & 5. (C)** Polar flagellar gene cluster and offside flagellar genes. The thick arrows represent the FlrA/RpoN- and FliA-dependent promoters known from *S. oneidensis* [14]. New identified σ^70^-dependent promoters are shown with thin arrows. The numbers indicate the base position in the chromosome.

The regulation of flagellar synthesis has been extensively studied for some Gram-negative gammaproteobacteria. At the transcriptional level, a three-tiered regulation cascade has been well-established for *Salmonella enterica*, which leads to a stepwise production and assembly of the flagellar multiprotein complex [11]. In polarly flagellated gammaproteobacteria, such as *Vibrio* sp. or *Pseudomonas aeruginosa*, a four-tiered transcriptional hierarchy underlies expression, production and assembly of the flagellar apparatus [12,13]. A variation of the four-tiered transcriptional cascade was shown to exist in the monopolarly flagellated gammaproteobacterium *Shewanella oneidensis* [14–16]. In this and other *Shewanella* species, the vast majority of flagella-related genes are clustered in several adjacent operons in the genome [14] (**Supplementary Fig. 1B**). Flagellar gene expression is initiated with the activation of the flagellar master regulator FlrA. In concert with sigma factor 54 (σ^54^; RpoN), FlrA activates expression of a number of genes encoding most building blocks of the cell envelope-embedded basal body and the hook. In addition, FlrA/σ^54^ induces expression of several further regulators, among which is the two-component system FlrBC. In polar flagellates such as *Campylobacter* (here designated FlgSR), *Vibrio* or *Pseudomonas* (here designated FleSR), these two-component regulators are thought to form a checkpoint that monitors MS-ring, C-ring and fT3SS assembly to then activate the genes encoding the building blocks for the flagellar rod and hook [12,13,17–19]. In contrast, in *S. oneidensis* these genes are under direct control of FlrA/σ^54^. In *Shewanella*, FlrBC appears to be generally dispensable for flagella synthesis, but may be involved in regulating the composition, mechanical properties and behavior of the flagellar filament [15,16,20,21]. Among the further regulators thought to be under direct control of FlrA is the sigma factor σ^28^ (FliA), which is responsible for expression of the late flagella factors, i.e., some motor components and the main flagellin. FliA is bound and thereby inactivated by its anti-sigma factor FlgM to prevent production of large amounts of flagellin before assembly of the filament can start. Only upon completion of the hook structure, FlgM is excreted from the cell and the released σ^28^ initiates expression of its corresponding promoters [11].

In polarly flagellated gammaproteobacteria, flagella synthesis needs to be spatiotemporally regulated to target assembly to the designated cell pole. In addition, the cells require a counting mechanism that restricts the number of flagella. In many bacterial species, proper targeting and counting is mediated by two proteins, FlhF and FlhG, which, in *Shewanella*, are also proposed to reside in the second tier of the transcriptional hierarchy under control of FlrA [14–16]. In the absence of FlhF, an SRP-type GTPase, less cells are flagellated and the flagellum is frequently detached from the cell pole [22–24]. The MinD-like ATPase FlhG antagonizes the function of FlhF at the cell pole and plays a major role in the counting mechanism that restricts the number of polar flagella to one [22,24–26]. Our previous work indicated that, in *Shewanella putrefaciens*, FlhG binds to the C-ring building block FliM as a monomer and passively travels to the nascent flagellar basal body. Assembly of FliM into the C-ring leads to release of FlhG and enables its ATP-dependent dimerization. As an ATP-bound dimer, FlhG directly interacts with FlrA. Thereby, FlhG prevents transcription of genes encoding further early basal body building blocks and thus effectively links C-ring and basal-body assembly with transcriptional control to limit the number of flagella. FlrA stimulates the ATPase activity of FlhG, which leads to monomerization of this regulator [25,26]. A similar spationumerical regulation mechanism has been shown to be active in *P. aeruginosa* with the FlhG and FlrA orthologs FleN and FleQ [27].

During our investigations on the interplay between FlhG and FlrA, we noticed that in mutants in which *flrA* was deleted, FlhG was present at levels similar to those of wild-type cells [26], which did not concur with the current models stating that FlhG expression and production is under control of FlrA. This prompted us to revisit the current flagella regulation model by determination of mRNA and protein levels in *S. putrefaciens* wild-type cells and mutants lacking the main flagellar regulators FlrA, RpoN (σ^54^) or FliA (σ^28^). We found that the level of a number of early basal body proteins and the regulators FlhF, FlhG and FliA remains unchanged in the absence of FlrA or RpoN, and we identified several constitutive promoters driving the transcription of genes encoding these components. We found that the constant level of FlhG is required to maintain the numerical control of flagella synthesis, demonstrating a significant physiological role of the constant production. Our results indicate that only some key components, which are required to initiate flagellar assembly, are strictly controlled at both the transcriptional and protein level, adding another level of complexity underlying the regulation of flagella synthesis.

## Results

### Determining the regulons of RpoN, FlrA and FliA in *S. putrefaciens*

The presence of FlhG in the absence of FlrA [26] lead us to speculate that the flagellar master regulator FlrA may not or only partially regulate expression of flagellar genes and presence of the corresponding encoded proteins. In addition, little data exists on the regulation of flagellar genes by FlrA in *Shewanella*, and, so far, the presence and position of most potential FlrA-dependent promoters relies on *in silico* predictions in *S. oneidensis*. As in the latter species, most genes required for the formation of the main polar flagellum of *S. putrefaciens* are clustered in one genomic region, and the gene organization closely resembles that of *S. oneidensis* [14,28] (**Supplementary Figure S1**). Further genes encoding proteins directly related to flagellar synthesis and function, the stator proteins PomA and PomB, the T-ring proteins MotX and MotY, and the motor switch-effector ZomB, are scattered throughout the chromosome [28,29]. In addition, *S. putrefaciens* harbors a secondary flagellar system, whose genes are clustered as a single operon elsewhere on the chromosome and which has not apparent role in the regulation of primary polar system [28].

To generate a comprehensive data set on the role of the main flagellar regulators in *S. putrefaciens*, we determined both mRNA and protein levels in mutant strains in which *flrA* (Δ*flrA*), *rpoN* (Δ*rpoN*) or *fliA* (Δ*fliA*) were removed from the genome by *in-frame* deletion. Earlier studies indicated that loss of the two-component system FlrBC, which in other species, such as *Vibrio cholerae*, is crucial for flagella formation [13], has no effect on flagella synthesis in *S. putrefaciens* under our standard experimental conditions [21]. To confirm this, flagella-mediated swimming of Δ*flrBC* mutants was determined and compared to that of wild-type cells and mutants bearing mutations in other main transcriptional regulators (Δ*flrA*; Δ*fliA*; Δ*flrABC*). While in the latter mutants spreading in soft agar was greatly diminished or completely abolished as expected, the Δ*flrBC* mutant showed no difference to wild-type spreading (**Supplementary Figure S2A**). Therefore, we concluded that the FlrBC two-component system has a similar role as in *S. oneidensis* and hence did not further consider *flrBC* in this approach.

RNA and protein levels of Δ*flrA*-, Δ*rpoN* and Δ*fliA*-mutant cells were determined during early exponential growth phase and compared to those of wild-type cells cultivated under the same conditions. As expected, the largest change occurred in the Δ*rpoN* mutant compared to the WT with 263 higher and 285 lower gene transcript levels. 104 genes were differentially regulated in the Δ*flrA* mutant among which 50 had higher and 54 had lower RNA levels (**Supplementary Table S1**). In the absence of *fliA*, 102 genes were expressed at higher and 80 genes at lower levels (**Supplementary Table S1**). As expected, many of the corresponding gene products have documented roles in flagella synthesis, but also a number of different genes not apparently related to flagella-mediated motility are directly or indirectly regulated by RpoN, FlrA and FliA at the transcriptional level.

At the protein level, FlrA-, RpoN or FliA-dependent changes were much less pronounced. The mass spectrometry approach identified 71 proteins in Δ*rpoN* mutants with significantly different abundances, 43 proteins in Δ*flrA* mutants, and 35 proteins in Δ*fliA* mutants compared to the wild-type proteome (**Supplementary Table S2**). Changes at the protein levels were reflected at the transcriptome levels, as genes encoding proteins that exhibited higher or lower abundance were similarly up- or downregulated in the corresponding mutants. For further analysis, we concentrated on those gene products that were directly or indirectly affected by RpoN, FlrA or FliA at the level of both RNA and protein (see **Supplementary Figure S3**).

For FlrA, the vast majority of these gene products were either directly associated with the primary flagellar system or the Bpf surface adhesion system, which has previously been found to be regulated by FlrA in *S. putrefaciens* [30]. Notably, FlrA appears to regulate the abundance of the flagellins FlaAB_2_ of the secondary flagellar system, indicating some cross talk between the two systems. Besides the flagellar components and the Bpf surface adhesion system, some further genes and gene products appear to be regulated by RpoN, FlrA and FliA (see **Supplementary Tables S1 & S2**). These apparent flagella-unrelated genes and proteins were not considered further in this study.

### Several flagellar proteins are present in the absence of FlrA, RpoN and FliA

The combined transcriptome and proteome analysis strongly hinted at the expected role of FlrA as a major flagellar regulator. However, expression and protein levels of several flagellar genes and the corresponding gene products did not exhibit major differences in the absence of *flrA*. As some flagellar proteins were not detected at sufficient reliability in the global mass spectrometry approach, the results were validated by Western immunoblotting. To enable specific detection of the appropriate proteins, strains were constructed in which the corresponding genes were extended on the chromosome by a sequence adding a C- or N-terminal 3xFLAG-tag as a suitable epitope to the produced protein. This way, we labeled FliE, FliF, FliG, FliH, FliI, FliJ, FliK, FliL, FliM, FliN, FliP, FliQ, FlhA, FlhB, FlhF, FlhG, FliA, FlrB, FlrC, FlgK, FlgM and MotY. For detection of FliO, the protein was fused C-terminally to sfGFP and for MotX a suitable N-terminal mCherry fusion was already established. Most labeled proteins were stably produced and supported flagella-mediated motility to almost wild-type levels (**Supplementary Figures S2B, S4 & S5**). Exceptions were the FLAG-tagged versions of FliE, FliK, FliN, FliQ and FlgK, which did not lead to functional flagella synthesis and the FliG- and FliP-FLAG mutants that were negatively affected in spreading through soft agar. Stable production still allowed to determine general differences in the presence of these proteins. To determine the presence or absence of the proteins in the presence of FlrA, RpoN or FliA, the tagged variants were introduced into *S. putrefaciens* strains lacking *flrA, rpoN* or *fliA* (**Figure 1B**).

Taken together, the global transcriptional and proteome data and immunoblotting approach indicate that several genes are transcribed and the corresponding proteins produced in the absence of FlrA or FliA (summarized in **Figures 1A** and **1C**). These comprise a whole gene cluster, *fliLMNOPQRflhB*, which likely forms an operon and encodes proteins required for the formation of part of the cytoplasmic C-ring and the ft3SS. Other constitutively produced proteins are further components of the ft3SS, FliI and FliJ. Also normally produced were the regulators FlhF, FlhG (as previously identified) and the σ^28^ factor FliA, which are thought to constitute a transcriptional unit. Similarly present are the anti sigma factor for σ^28^, FlgM, along with cytoplasmic export chaperone FlgN. Also here both corresponding genes likely form an operon. Surprisingly, also the motor components PomA and PomB, the flagellar stators, and the T-ring protein MotX, which is required for stable recruitment of the stators into the flagellar motor, are constantly produced. Almost all other components showed a pronounced downregulation at the transcriptional level upon deletion of *flrA* or *rpoN*, and the corresponding proteins could not be detected. Among these genes and gene products are some components required for initiation of basal body formation, e.g., the ft3SS component FlhA, and the MS- and C-ring proteins FliF and FliG, and virtually all components of the rod, the outer rings, the hook and the flagellins. As previously predicted, σ^28^ (FliA) was found to be crucial for the expression and production of the main flagellin FlaB_1_, FlaG, and, in addition, also for the lacking T-ring component MotY.

### Identification of constitutive promoters driving expression of flagellar genes

Our results demonstrated that the presence of a set of flagellar proteins is not dependent on FlrA. This indicated that the corresponding gene expression may occur even when *flrA* is deleted. We therefore determined if additional FlrA-independent promoters may drive expression and production of at least some flagellar genes and proteins. To this end, a 400 to 550 bp DNA region upstream of selected flagellar genes, which putatively contained such FlrA-independent promoter regions, were amplified and fused to the *Photorhabdus luminescence luxCDABE* operon on a broad-host range plasmid. Based on the occurrence of proteins and the predicted operon structure (**Figure 1C**), we selected regions upstream of the following genes: *fliF, fliG, fliH, fliI, fliK, fliL, flhA, flhF, flhG* and *fliA*. The region upstream of *fliE*, bearing a highly conserved FlrA-binding site, was used as a positive control. The plasmids were transformed into *S. putrefaciens* wild-type, Δ*flrA* and Δ*flrABC* cells, and emission of light due to expression of the *lux* genes was measured in the presence and absence of FlrA (or FlrABC) (**Figure 2A; Supplementary Figure S6A**). As expected, the positive control bearing the upstream region of *fliE* displayed a strict FlrA-dependent expression. A similar pattern occurred for *fliK* and *flhA*, suggesting that expression of the two genes is predominantly or exclusively governed by FlrA. In sharp contrast, for the *fliF, fliG, fliL, flhF, flhG* upstream regions *lux* expression occurred both in the presence and absence of FlrA. According to the determined light emission, the highest promoter activity is located in front of *fliL*. Also the *fliA* upstream region enabled *lux* expression, albeit at a lower level. For *fliF, fliG, flhG* and *fliA*, there was no significant difference in expression in both the wild-type and Δ*flrA* strain. Expression initiated from the upstream regions of *fliL* and *flhF* was lower in the absence of FlrA, however, still robust luminescence occurred. These results strongly suggested that within these regions, FlrA-independent promoter regions are present that drive constitutive expression of the downstream regions. No promoter activities were detected for the *fliH* and *fliI* genes (**Supplementary Figure S6B**).

**Figure 2:**
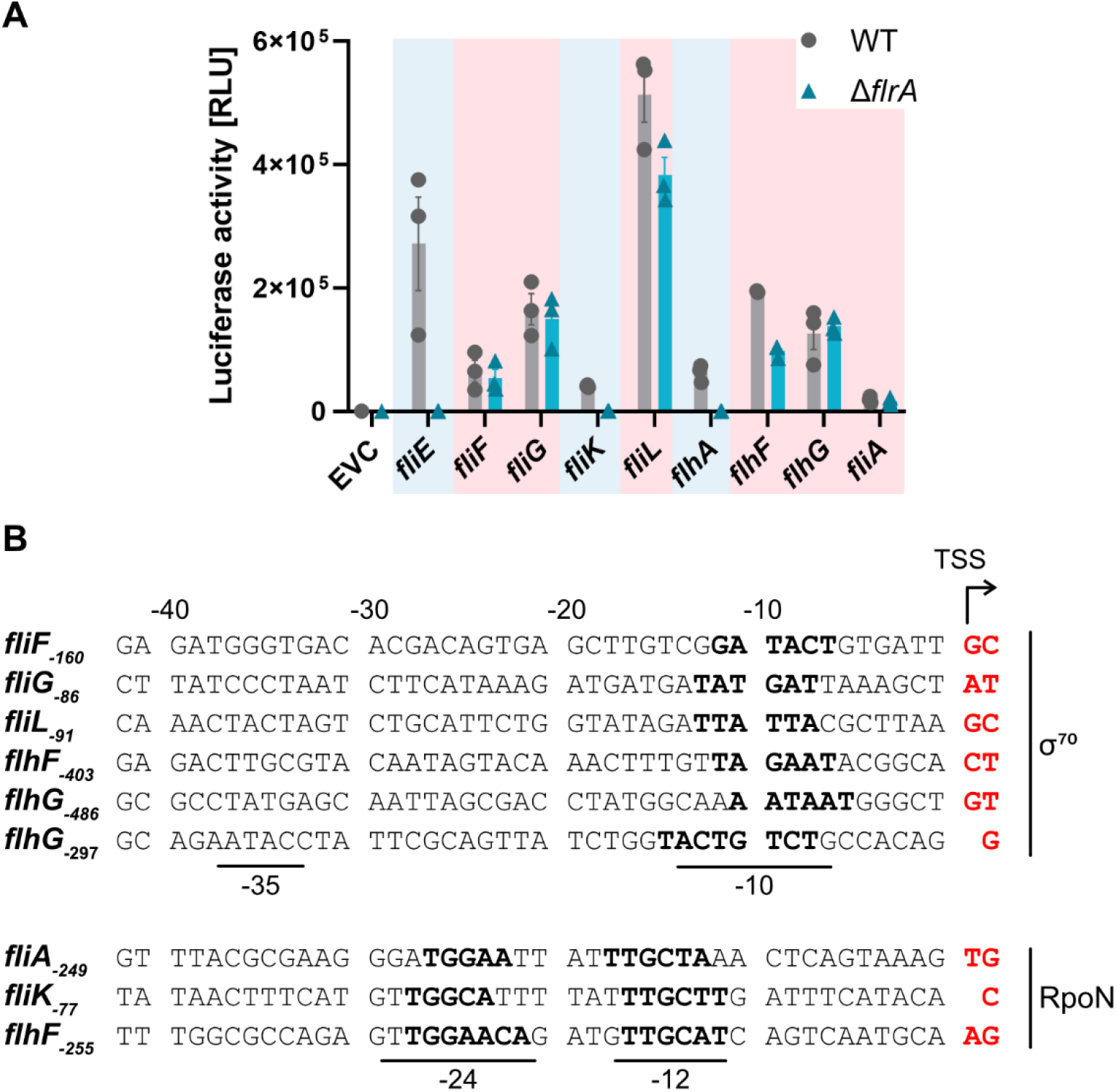
Some promoters of early flagellar components are not dependent on FlrA. **(A)** Determination of the promoters’ transcriptional activity using a luciferase reporter system. The 400 or 550 bp fragments upstream of the translational start were fused to the reporter genes *luxCDABE* from *Photorhabdus luminescence*. These reporter plasmids with the promoter regions of interest were analyzed in *S. putrefaciens* WT and deletion of the master regulator *flrA* (Δ*flrA*). The promoters highlighted in blue are FlrA-dependent and those highlighted in red are active in the absence of FlrA. Shown are the means and standard deviations from biological and technical triplicates. **(B)** Identification of transcription start sites (TSS) by 5’ RACE-PCR. Bases highlighted in red and the curved arrow indicate the TSS. In the DNA sequence upstream from the TSS, the binding motifs of the transcription factors of σ^70^ (−10 and -35 region) and RpoN (−12 and -24 region) were highlighted in bold. The numbers at the genes indicate the base pairs for the distance from the TSS to the translational start.

To identify the corresponding promoter region, the promoter-containing DNA fragments were truncated until loss of expression occurred to roughly home in on the position of the promoters (**Supplementary Figure S6B**). Then, we performed ‚Rapid Amplification of 5’ cDNA-Ends’ (RACE) PCR experiments using the *lux*-based reporter plasmids as templates in *S. putrefaciens*. By this, several putative transcriptional start sites were identified and mapped upstream of *fliF* (160 bp; *fliF*_-160_), *fliG*_- 86_, *fliK*_-77_, *fliL*_-91_ and *fliA*_-249_ (**Figure 2B**). Two potential transcriptional start sites occurred upstream of *flhF* (*flhF*_-403_; *flhF*_-255_) and flhG (*flhG*_-486_; *flhG*_-297_). Based on consensus sequence predictions, the deduced -10 and -35 regions of *fliF*_-160_, *fliG*_-86_, *fliL*_-91_, *flhF*_-403_, and both *flhG* putative transcriptional start sites (*flhG*_-486_ and *flhG*_-297_) may represent (degenerated) σ^70^-dependent promoters. As σ^70^ is the ‚house keeping’ sigma factor of *S. putrefaciens*, this would explain constant expression of the downstream genes. Homologies of the deduced -10 and -24 regions upstream of *fliA*_-249_, *fliK*_-77_ and *flhF*_-255_ to the binding sequence of RpoN (σ^54^) indicated that these potential promoters may be driven by RpoN, the main sigma factor involved in flagella synthesis. For an overview of the newly mapped promoters’ position, see **Figure 1C**.

### Constitutive production of FlhG is required for proper numerical control of flagella synthesis

The presence of a number of constitutively active promoters within the operons encoding part of the flagellar basal body as well as the spatial and numerical regulators FlhG and FlhF and the sigma factor enabling production of the main flagellin FlaB_1_, FliA, suggested that constant protein production may have a role in flagellar synthesis. This particularly applied to FlhF and FlhG as changes in the levels of these proteins affect the number of the polar flagella in *S. putrefaciens* [25,26]. To determine a potential effect on flagellar synthesis, we aimed at silencing the promoters on the chromosome in the wild-type background. This was complicated by the fact that the identified promoters exhibit poor consensus sequence conservation and were, additionally, often located within the upstream structural gene of the same operon. We therefore introduced base substitutions within the predicted operons sequences without changing the encoded protein’s codons or disturbing the open reading frame of the corresponding gene (**Supplementary Table S3**). An effect of the base substitutions on transcriptional activity was first determined using the *lux*-based reporter systems (Fig. 3A). By this, we were unable to generate substitution mutants with a notable effect for the promoter upstream of *fliL*. In contrast, we successfully silenced the promoter upstream of *flhF* within the gene *flhA* (P1; **Figure 3A**), another promoter upstream of the two promoters within *flhF* (P2) and diminish the activity of the third, more downstream (P3), both in front of *flhG*. These mutations were then introduced into their native positions within the chromosome. By western blotting we determined the protein levels of FlhF and FlhG (**Figure 3B**), and found that FlhG was produced at a slightly lower level in the mutants lacking the three promoters. Only in the additional absence of FlrA, the protein levels were below the detection limit. Notably, the latter also occurred for FlhF, indicating that both proteins are controlled by the master regulator FlrA. The strain with the three substituted promoters was then tested on its ability to assemble flagella and to spread through soft agar. Despite the production and presence of residual FlhG levels, the mutant exhibited a significant phenotype: while the number of flagellated cells was highly similar in wild-type and mutant cells (about 75%), about 25% of the mutants cells synthesized more than one filament at the cell pole, which was never observed for the wild-type (**Figure 3C, D**). Also the spreading capability of the mutant cells through soft agar was significantly decreased (**Figure 3E**).

**Figure 3:**
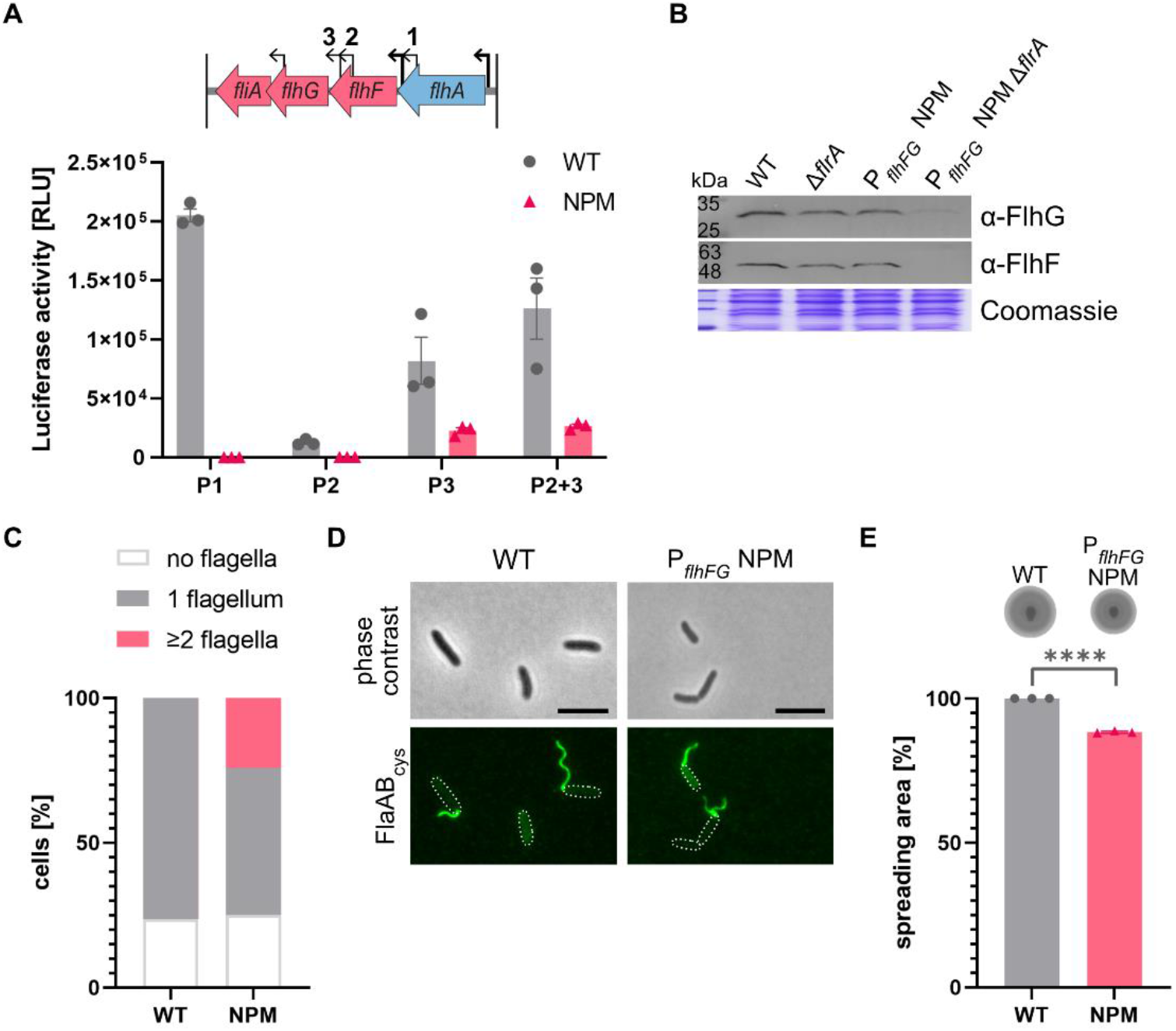
Constitutive production of the flagellation regulators FlhF and FlhG are essential for monopolar flagellation. **(A)** Top, overview of the *flhA* operon is shown with regulation of gene expression (blue: FlrA/RpoN-dependent, red: present in Δ*flrA*, Δ*rpoN*, Δ*fliA*) and all promoters (thick arrow: FlrA/RpoN-dependent promoter, thin arrow: σ^70^-dependent promoter). Below are the measurements of promoter activities using the luciferase reporter system. The codon usage of the sequences of the σ^70^-dependent promoters of *flhF* (P1) and *flhG* (P2, P3, P2+3) were substituted (NPM, no promoter motif) to obtain the lowest possible promoter activity and compared with the native promoter sequence (WT). Shown are the means and standard deviations from biological and technical triplicates. **(B)** Western blot to check the protein levels of FlhF and FlhG in backgrounds of *flrA* deletion (Δ*flrA*), low σ^70^ promoter activity of *flhFG* (P_flhFG_ NPM) genes, and the P_flhFG_ NPM Δ*flrA* combination. Coomassie stained SDS-polyacrylamide gel serves as loading control. The complete blots are shown in **Supplementary Figure 5D. (C)** Quantification of the number of flagella per cell. Flagellins were labeled via cysteine exchange using the fluorescent dye maleimide-CF484. Quantification was performed using 400 cells in biological triplicates. **(D)** Microscopy images of flagellar staining of the P_flhFG_ NPM strain compared with the WT. The scale represent 5 μm. **(E)** Top, the spreading of the *S. putrefaciens* WT and chromosomal mutant NPM (P_flhFG_ NPM) showing lowest possible promoter activity of all three σ^70^- promoters of *flhFG* are shown on soft agar plates. Below is the percent spreading radius from these two strains quantified. The percent spreading radius of P_flhFG_ NPM was normalized to the WT. The experiment was performed in biological triplicates. Asterisks indicate that the difference from the WT is significant with a *p*-value of <0.0001 (two-sample t-test).

Taken together, the results strongly indicate that expression from constitutive promoters and production of FlhG and maybe FlhF is required to maintain its role as a numerical regulator of flagellar synthesis.

### RpoN and FliA are involved in regulation of the Bpf surface adhesion system

In both RNA and proteome analyses the gene cluster and the gene products of the *bpf* operon, which encodes a surface adhesion protein system, showed higher abundance upon deletion of *flrA*. This finding was in accordance with a previous study reporting that, in addition to its role as a flagellar master regulator, FlrA serves as transcription repressor of the *bpf* operon [30]. In addition to FlrA, also loss of RpoN and FliA resulted in significantly different Bpf RNA and protein levels (**Figure 4A**). However, in contrast to FlrA, RpoN and FliA rather appears to positively regulate expression of *bpf* genes and corresponding gene products, as both show a lower abundance in the Δ*rpoN*, Δ*fliA* mutant background. To confirm this, the genes encoding two major proteins of the system, BpfA and AggA, were mutated on the chromosome to result in the production of C-terminally FLAG-tagged versions of the protein. Both proteins showed a higher abundance in the absence of FlrA and a lower abundance in Δ*rpoN* and Δ*fliA* mutants (**Figure 4B**). Thus, FliA is directly or indirectly required for normal expression of the Bpf surface adhesion system in *S. putrefaciens*.

**Figure 4:**
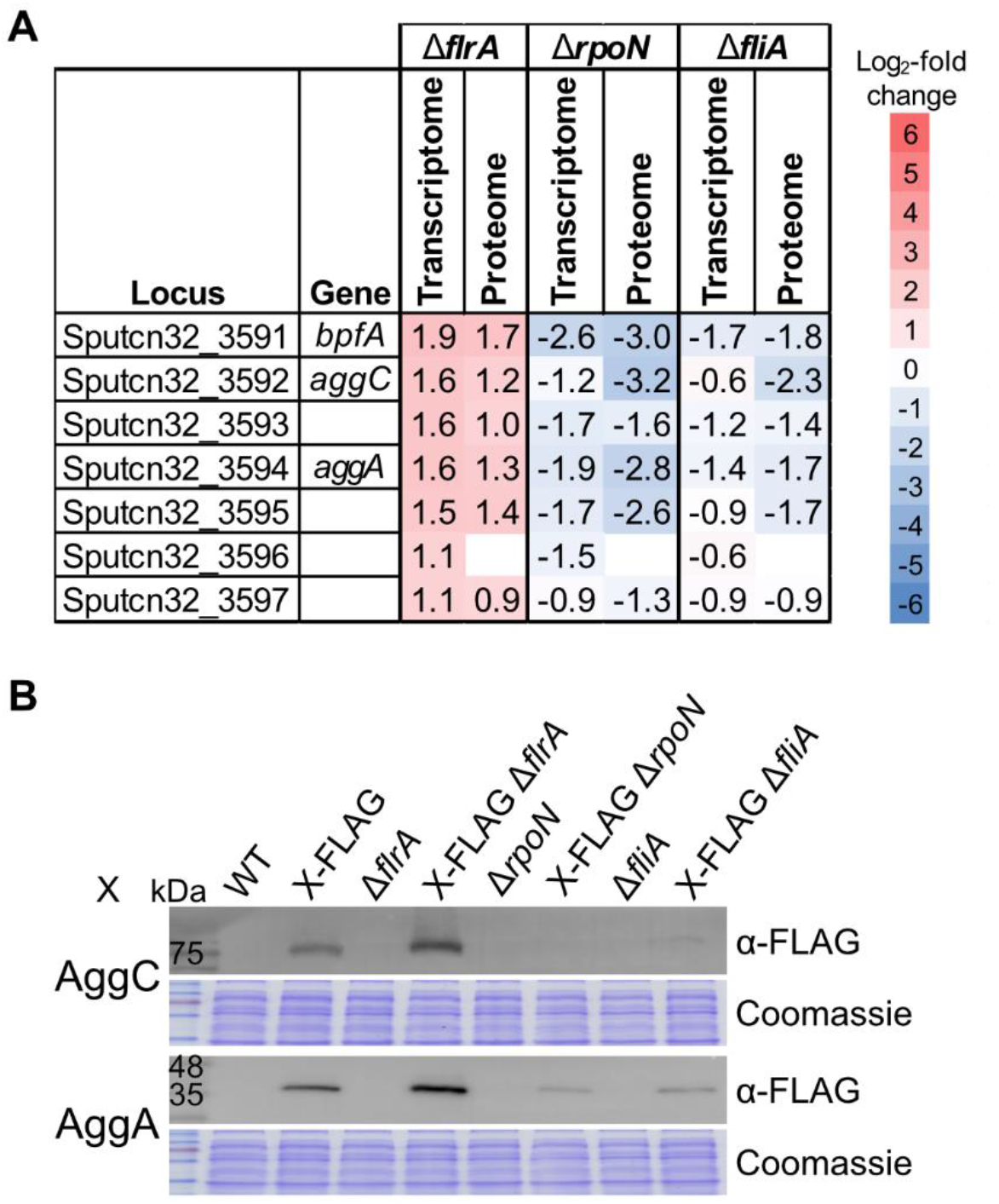
Production of the Bpf surface adhesion system is inhibited by FlrA and activated by RpoN and FliA. **(A)** Analysis of the deletion strains *flrA, rpoN*, and *fliA* by RNA sequencing (transcriptome) and mass spectrometry (proteome). The log_2_-fold changes in mRNA or protein levels of the deletion strains compared with *S. putrefaciens* WT are indicated. **(B)** Western blot analyses of components of the Bpf system AggC-FLAG and AggA-FLAG in *S. putrefaciens* deletion strains Δ*flrA*, Δ*rpoN*, or Δ*fliA*. Coomassie-stained SDS-polyacrylamide gel serves as loading control. The complete blots are shown in **Supplementary Figure 5C**.

## Discussion

In this study, we have used complementary transcriptomic and proteomic approaches to investigate the regulatory network underlying the synthesis of the single polar flagellum in *S. putrefaciens*. Particularly, the quantification of protein levels allowed us to identify flagellar components and regulators whose production is strictly controlled. Jointly, the results demonstrate that the *S. putrefaciens* flagellar regulation significantly deviates from the four-tiered regulatory pathway model that is commonly proposed for monopolarly flagellated species such as *Vibrio* or *Pseudomonas* [12,13,31]. We show that, under standard conditions used in the experiments, a number of flagellar proteins including those forming the chemotaxis systems are constitutively produced (see **Figure 1**), and that production of just three proteins directly related to flagellar synthesis, FlaB, FlaG and MotY, is controlled by FliA (σ^28^). All residual flagellar proteins depend on FlrA/RpoN, while the regulators FlrB and FlrC are not required. Taken together, the data indicates that, at least under these conditions, *S. putrefaciens* rather possesses a three-tiered flagellar transcriptional regulatory pathway (**Figure 5**). We propose that the top-tier regulator FlrA (in concert with σ^54^ (RpoN)) activates expression and production of numerous flagellar building blocks, which, together with the already present constitutively produced components result in localized assembly of the flagellar basal body, the rod and the hook. Completion of the hook then allows export of the anti-sigma factor FlgM to release σ^28^ (FliA) [11]. Free σ^28^ in turn enables initiation of synthesis of the major flagellin FlaB and the T-ring component MotY. While FlaB is being produced, the already synthesized minor flagellin FlaA can be exported and assembled to form the cell proximal part of the flagellar filament. This results in the typical spatial distribution of flagellins in the *S. putrefaciens* filament that optimizes its stability for a range of environmental conditions [21]. Together with MotX, MotY assembles the periplasmic T-ring, which allows functional coupling of the already produced PomAB stators into the now completed flagellar motor, which can start the rotational movement of the still growing flagellum. The function of the third σ^28^-controlled flagellar protein FlaG is yet elusive. Based on the findings in other species, FlaG may have a role in regulating the length of the filament [32–34]. Thus, the transcriptional hierarchy of regulation is rather limited in *S. putrefaciens*. This strongly indicates that flagellar assembly can generally occur also in the absence of extensive transcriptional regulation, similar to that of the highly homologous type III secretion system [35,36].

**Figure 5:**
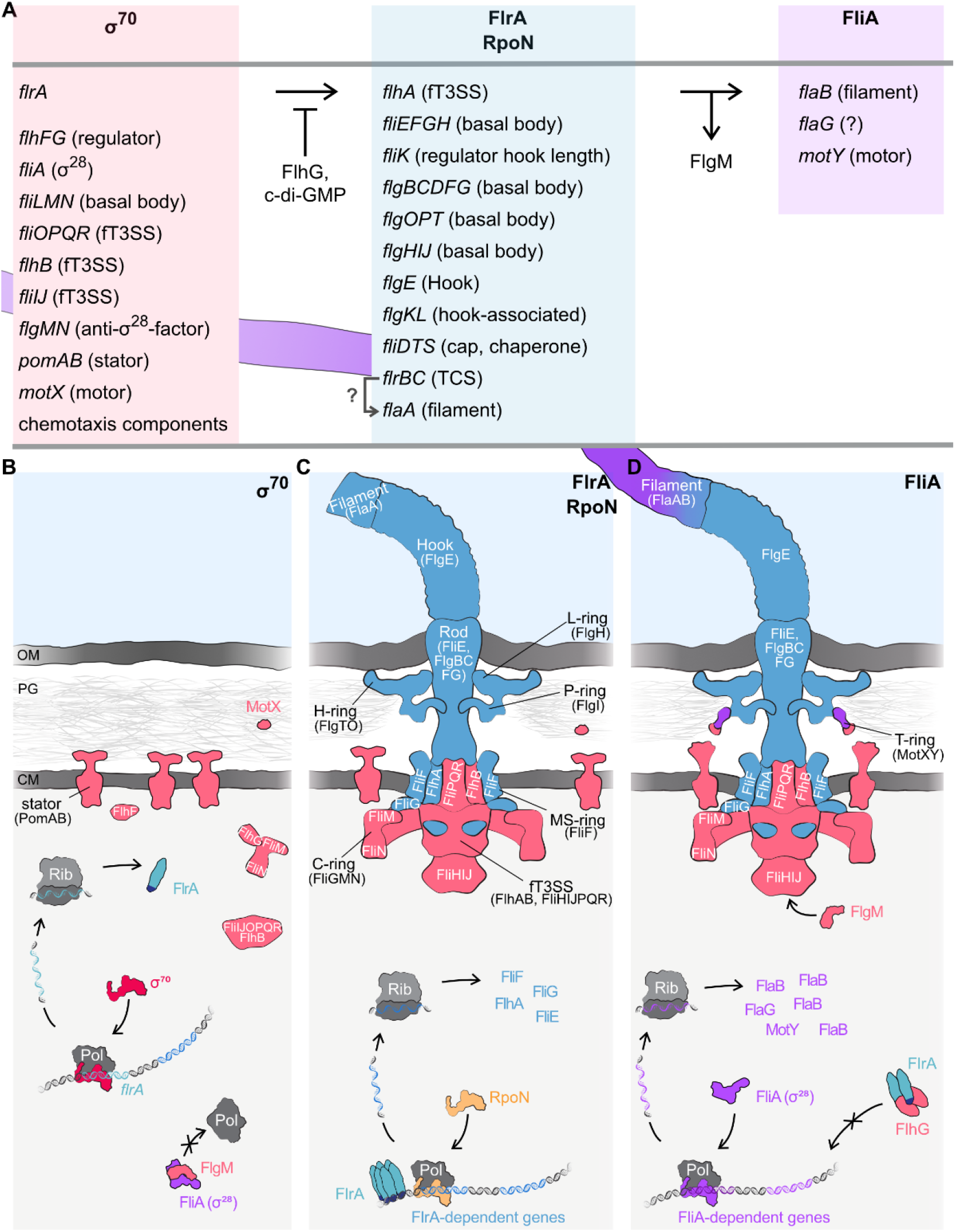
Transcriptional regulation model of polar flagellar synthesis in *S. putrefaciens*. **(A)** Transcriptional hierarchical regulation. **(B)-(D)** Schematic overview of flagellar assembly. Figures have been modified from [21]. The master regulator flrA as well as other flagellar regulators (*flhFG, fliA, flgM*) and components of the basal body are σ^70^-dependent. Binding of FlhG or c-di-GMP to FlrA represses transcriptional activity. FlrA/RpoN activate the expression of additional genes. These include components of the basal body, hook, and first component of the filament. After assembly of the hook, the anti-sigma factor FlgM is exported and FliA can activate gene expression of late flagellar genes. These include the major flagellin *flaB* as well as the motor protein *motY*. Flagellar components labeled in red are dependent on the housekeeping factor σ^70^. Gene expression of components labeled in blue are activated by FlrA as well as RpoN and components labeled in purple are activated by FliA. For further explanations see the Discussion section.

A notable feature of the less intricate flagellar transcriptional hierarchy in *S. putrefaciens* is the missing requirement of the two-component system FlrBC, which has previously been demonstrated also for the closely related species *S. oneidensis* [15,16]. In *V. cholerae* and *P. aeruginosa*, the corresponding orthologs (FlrBC in *Vibrio* and FleSR in *Pseudomonas*) are crucial for a production of a significant subset of flagellar proteins, in particular those for synthesis of the flagellar rod and hook structures. Accordingly, no functional flagella can be synthesized in the absence of these regulators in these species [12,37–40]. It has been proposed that, as shown for the *Campylobacter jejuni* FlgSR two component system, FlrBC and FleSR form a regulatory checkpoint for completion of the MS-ring, the C-ring and the ft3SS, before assembly of the following parts is initiated [19]. This checkpoint appears to be absent in *Shewanella* without negatively affecting flagellar synthesis. In *S. oneidensis*, the FlrBC is thought to be involved in regulating abundance of the minor flagellin FlaA, which may influence the stiffness and geometry of the flagellar filament and, by this, the swimming performance [15,16], which could not be observed in *S. putrefaciens* [21]. It is possible that *S. putrefaciens* FlrBC is activated and has a regulatory function under particular environmental conditions, but this remains to be shown.

An unexpected finding of our study was the number of flagellar proteins that were present at normal abundance also in the absence of FlrA/RpoN. Among these proteins were most, but not all, components of the ft3SS and two of the three building blocks forming the cytoplasmic C-ring (FliM and FliN). Also three important regulators are among the constitutively produced flagellar proteins, FlhF, FlhG, and the sigma factor FliA (σ^28^) along with its anti-sigma factor FlgM. For most of the corresponding genes or gene clusters, we could identify appropriate promoters. One possible explanation is that strict regulation of early building blocks has been lost during evolution except for some key elements, such as FliF, FliG and FlhA, to initiate formation of the MS- and C-ring, and the ft3SS, respectively. This would, however, occur at the cost of the energy required for the constant protein production. In addition, the number of identified promoters that are particularly present upstream of the regulator-encoding genes *flhF, flhG* and fliA rather pointed at a biological function of the constitutive production. In this study, we could identify at least one of such possible functions, which is related to the regulator pair FlhF and FlhG.

The SRP-type GTPase FlhF and the MinD-like ATPase FlhG are implicated in flagellar positioning and number in a number of bacterial species [9,24,41]. As in most other polarly flagellated species where FlhF has been studied, the latter is a positive regulator of flagellar synthesis and serves as a determinant for polar localization of the flagellum in *Shewanella* [22,23]. In *S. putrefaciens*, deletion of *flhF* results in a lower number of flagellated cells with frequent flagellar mi**s**placement away from the cell pole, while overproduction of FlhF gives rise to a polar flagellar bundle [23]. In contrast, loss of *flhG* leads to polar hyperflagellation, while *flhG* overexpression significantly decreases flagellar formation [25,26]. The current model for polarly flagellated bacteria suggests that an ATP-bound dimeric FlhF localizes to the cell pole and recruits FliF and/or facilitates assembly of the MS-ring as the earliest component of the bacterial flagellum in the cytoplasmic membrane [24,42]. In *S. putrefaciens*, monomeric FlhG binds to the C-ring component FliM(N) and the complex travels to the nascent basal body [25]. Upon assembly of FliM into the C-ring, FlhG is released, and, as a membrane-associated ATP-bound dimer, stimulates the GTPase activity of FlhF [23,43], resulting in FlhF monomerization and loss of polar localization. In addition, the ATP-bound FlhG dimer can interact with FlrA, which interferes with FlrA activity as an activator of the early flagellar transcription, thereby shutting down synthesis of more earlier building blocks upon completion of C-ring assembly [26]. Notably, FlrA stimulates FlhG ATPase activity which results in FlhG monomerization and loss of FlrA interaction. Accordingly, interference with any of these protein interactions or the protein ratios results in at least partial loss of the proper regulation of flagellar counting and placement [26]. To keep the number of polar flagella to one, the regulating partner switch has to be maintained also after the flagellar synthesis has been completed to prevent the initiation of a next round of flagellar formation. We have shown here that partial silencing of the promoters upstream of *flhG* result in cells with more than a single polar filament. We therefore propose that constitutive FlhG production of helps to maintain a level of protein that balances initiation and prevention of forming additional flagella and thus renders the flagellar counter more robust.

So far, we were not able to sufficiently modify the predicted promoter regions in a way to suppress permanent expression of the other regulator-encoding genes to address if this has any effect on flagellar synthesis and activity. Given the roles of FlhF in the flagellar placement and counting regulation, such a role is easily conceivable. Notably, also FliM(N), the C-ring-binding partners of FlhG, are among the constitutively produced proteins. It has been shown for *Escherichia coli* flagellar motors that FliMN are constantly exchanged in the rotating flagellar motor [44,45]. Thus, we hypothesize that a similar exchange in the *S. putrefaciens* motor would promote polar FlhG trafficking to maintain the regulatory circle as described, however, this remains speculative so far. A reason for FliA (σ^28^) being constitutively produced (along with its anti-sigma factor FlgM) is not obvious as our results rather predict that the regulatory role of this sigma factors is, as expected, in initiating production of the main flagellin and enabling assembly of the T-ring to allow coupling of the stators to the motor to start rotation. Also the role of FliA and RpoN in regulating the activity of the Bpf adhesin, which we observed, remains obscure. Previous work provided evidence that FlrA serves a repressor for expression of the *S. putrefaciens bpf* operon [30], which we similarly observed in this study. By this, FlrA reciprocally (co-)regulates the switch between the sessile and the free-swimming lifestyle. However, having RpoN and FliA as direct or indirect positive regulators of both motility and surface association appears counterintuitive and further studies are required to solve the role of RpoN and FliA in this process.

Taken together, our study demonstrates another level of regulation underlying the usual transcriptional hierarchy proposed for flagellar synthesis. It is yet unclear, if this mode of regulation also applies to other bacteria. Transcriptional studies on different species identified genes that appear to be constitutively transcribed, e.g, the *fliHIJ* gene cluster in *Pseudomonas putida* [46] or *flgA* and *fliEFGHIJ* in *Vibrio campbellii* [47]. Also in *P. aeruginosa*, similar genes and operons as in *S. putrefaciens* (*fliLMNOPQR-flhB; fleN-fliA*) are only slightly regulated at the transcriptional level [12], suggesting that the proteins may be permanently present. Also a sigma 70-dependent promoter for *flhF* has been identified in *P. putida* [48]. However, while numerous studies have addressed the transcriptional hierarchy of flagellar synthesis in an array of different bacterial species, surprisingly little is known about the regulation of flagellar protein levels in the cells. A study comparing the proteome of *Xanthomonas oryzae* wild type to that of a FleQ (FlrA) mutant identified surprisingly few proteins related to flagella-mediated motility [49], indicating a significantly different regulation cascade in this species. We expect that more proteomic studies on flagellar regulation will uncover similar or additional principles of regulating flagellar synthesis and assembly. For monopolarly flagellated bacteria, such as *S. putrefaciens*, it is important that upon completion of the flagellar basal body, re-initiation of flagellar assembly of another flagellum is prevented, so the copy number of at least the key initial proteins, FliF, FliG, and FlhA, need to be sufficiently low to inhibit the start of another round of flagellar assembly. To this end, the production of these proteins could be extremely well controlled or the surplus of these key building blocks is rapidly cleared from the cell. How *S. putrefaciens* in particular and bacteria in common are achieving this is currently under investigation.

## Materials and Methods

### Bacterial strains, growth conditions and media

Strains of *Escherichia coli* and *Shewanella putrefaciens* CN-32 used in this study are listed in **Supplementary Table S3**. *Shewanella* strains were grown at room temperature or 30 °C in LB medium, *Escherichia coli* at 37 °C in LB medium. The Medium of the 2,6-diaminopimelic acid (DAP)-auxotroph *E. coli* WM3064 were supplemented with DAP to a final concentration of 300 µM. When appropriate, selective media were supplemented with 10 % (w/v) sucrose, 50 mg/ml kanamycin, 2 % (w/v) L-arabinose or 40 ng/ml anhydrotetracyclin (AHT). Soft agar plates were prepared with 0.25 % (w/v) select agar and LB.

### Vector and strain constructions

All plasmids and oligonucleotides used in this study are listed in **Supplementary Tables S4** and S**5**. Construction of plasmids was performed using Gibson assembly [50]. Generation of markerless *in-frame* deletions and chromosomal integrations of gene variants or fusions in *S. putrefaciens* were carried out using the suicide vector pNPTS 138-R6K with EcoRV as previously described [51]. In order to overproduce the flagellar main regulator FlrA (Sputcn32_2580) the vector pBTOK [23] or a chromosomal integration into the major arabinose degradation pathway as previously described [52] were used. For analysis of flagellar promoter activity in *S. putrefaciens*, a transcriptional terminator cassette and the *luxCDABE* operon of *Photorhabdus luminscence* were amplified from pBBR1-MCS5-TT-RBS-lux [53] and cloned into the vector pBBR1-MCS2, using SacI and KpnI, respectively, yielding vector pBBR1-MCS2-TT-RBS-lux. The promotor regions to be analyzed were inserted upstream of the lux operon using XbaI and BamHI. Vectors were introduced into the appropriate strains via conjugation using *E. coli* WM3064 as a donor.

### Immunoblot analysis

Regulation of protein amount were determined by western blot analysis. Collection of protein samples, protein separation and immunoblot detection were carried out as previous described [25,26,28].

### Quantitative RT-PCR

Total RNA of exponentially growing *Shewanella* cells were isolated as previous described [26]. Residual DNA was removed with the Turbo DNA-free Kit (Invitrogen) according to the manufacter’s protocol. For the quantitative Real-Time PCR (qPCR) the CFX Connect Real-Time System (Bio-Rad) and white strips of low profile tubes with ultra clear caps (Thermo Fisher Scientific) were used for PCR amplification. The qPCR were performed as previously described [26]. Oligonucleotides (Sigma-Aldrich) used for qPCR are listed in **Supplementary Table S5**. Primer efficiencies and relative transcript levels were calculated according to Pfaffl [54]. Ct values for each gene of interest were normalized against the Ct value of Sputcn32_2070 (*gyrA*) and computed relative to the respective control.

### Luciferase assay

Measuring transcriptional activities of promoters with fusions to the *luxCDABE* operon were performed as previously described [28]. *Shewanella* strains were diluted to an OD_600_ 0.02 from overnight culture and grown under aerobic conditions to an OD_600_ 0.5-0.6 at 30 °C. Exponential growing cells are transferred in 160 µl aliquots to a white 96-well polypropylene microliter plate (Greiner). Luminescence emission was measured in technical and biological triplicates using a Tecan Infinite M200 plate reader (Tecan). The relative luminescence units (RLU) were calculated by dividing the luminescence intensity by its corresponding OD_600_ value to normalize the luminescence to an OD_600_ 1.

### 5’ sRACE-PCR

For identification of transcriptional start sites (TSS) were 5’
sRACE PCR performed. DNA-free RNA of *Shewanella* strains with the promoters to be analyzed on the pBBR1-MCS2-TT-RBS-lux plasmid were isolated of exponentially growing cultures. The further procedure were performed by the 5′/3′RACE PCR Kit, 2^nd^ Generation (Roche) according to the manufacter’s protocol. The amplification of dA-tailed cDNA in a first and second PCR were performed with Biozym Taq DNA Polymerase (Biozym) according to the manufacter’s protocol. The corresponding oligonucleotides are listed in **Supplementary Table S5**. Ligation of 1 µl 5′RACE PCR products were performed with the Qiagen PCR cloning kit (Qiagen) according to the manufacters protocol. Inserts of plasmids were sequenced with oligonucleotide pUCM13-52 by Sanger sequencing (Microsynth seqlab).

### Flagellar staining and microscopy

Fluorescent staining of polar flagellar filaments (CN-32 FlaAB_cys_) were carried out as previously described [20,23,55] using CF488A maleimide (Sigma-Aldrich). Fluorescence images were recorded by a DMI6000 B inverse microscope (Leica) equipped with a pco.edge sCMOS camera (PCO) and an HC PL APO 100x/1.40 Oil PH3 phase contrast objective using the VisiView software (Visitron Systems GmbH). Images were further processed using the ImageJ-based Fiji tool [56] and Affinity Designer 1.7v (Serif).

### Motility assay

Spreading ability of *Shewanella* strains in semi-solid environments were analyzed by placing 2.5 µl of an exponentially grown culture on soft agar plate. After incubation for 16 h at room temperature the plates were scanned for documentation using an Epson Perfection V700 Photo Scanner (Epson).

### RNA sequencing and data analysis

Bacteria were grown in 10 ml LB to early stationary phase (OD600 0.5, 30°C, 180 rpm). RNA was extracted using the RNeasy Mini Kit (Qiagen) and Qiashredder columns (Qiagen) according to the manufacturer’s instruction. Quality of the obtained RNA was checked using the RNA Nano Kit on an Agilent Bioanalyzer 2100 (Agilent Technologies). The removal of ribosomal RNA was performed using the Ribo-Zero Bacteria Kit (Illumina) and cDNA libraries were generated with the ScriptSeq v2 Kit (Illumina). The samples were sequenced in single end mode on an Illumina HiSeq 2500 device. Sequencing was conducted on RNA obtained from two independent growth experiments. Raw reads were trimmed using the tool ‘cutadapt’ (version 3.5) [57] with customized settings (--nextseq-trim=20; -g ‘TTTTT;min_overlap=1’; -m 36). Mapping was performed with ‘bowtie2’ (version 2.3.5.1) [58] with the settings “--very-sensitive-local -I 100 -X 1200” and with the NC_009438.1 (*Shewanella putrefaciens* CN-32) genome as a reference. Reads per gene were extracted with the tool ‘featureCounts’ (version 2.0.1) [59] (**Supplementary Figure S7B**). Data analysis including the MDS plot was performed with the R package edgeR (v.3.32.1) [60] (Supplementary Figure 7A). qPCR was conducted on *fliF* and *fliM* as another control (**Supplementary Figure S7C**). Significant regulation was accepted at a log_2_ fold change of less than -1 or greater than 1 and a *p*-value of less than 0.05.

### Mass spectrometry

Proteomic analyses were performed as described recently [61]. In short, cell pellets were lyzed by 2% sodium laroyl sarcosinate (SLS) and heat. Following reduction and alkylation, 50 µg of extracted protein was digested by trypsin. Post digest the detergents were precipitated by adding acid and peptides were desalted using C18 solid phase extraction. 1 µg of peptides were then analyzed by liquid chromatography-mass spectrometry carried out on a Q-Exactive Plus instrument connected to an Ultimate 3000 RSLC nano. Peptide separation was performed on a reverse-phase HPLC column (75 µm x 42 cm) packed in-house with C18 resin (2.4 µm, Dr. Maisch). The following separating gradient was used: 98% solvent A (0.15% formic acid) and 2% solvent B (99.85 acetonitrile, 0.15% formic acid) to 25% solvent B over 105 minutes and to 35% solvent B for additional 35 minutes at a flow rate of 300 nl/min. The data acquisition mode was set to obtain one high resolution MS scan at a resolution of 70,000 full width at half maximum (at m/z 200) followed by MS/MS scans of the most intense ions. The dynamic exclusion duration was set to 30 seconds. The ion accumulation time was set to 50 ms for MS and 50 ms at 17,500 resolution for MS/MS. The automatic gain control was set to 3×10^6^ for MS survey scans and 1×10^5^ for MS/MS scans.

Label-free quantification (LFQ) of the data was performed using Progenesis QIP (Waters) and SafeQuant. The strategy has been further described in [62,63]. Significant regulation was accepted at a log_2_ fold change of less than -1 or greater than 1 and a *p*-value of less than 0.05.

## Acknowledgments

The authors thank Ulrike Ruppert for excellent technical support. The study was supported by a grant (TRR 174 P12) to KMT from the Deutsche Forschungsgemeinschaft within the framework of the Transregio program TRR 174. K.P. acknowledges funding by the Deutsche Forschungsgemeinschaft (EXC2051-ID390713860) and the Vallee Foundation.

## Data availability

The RNA seqeuncing data has been deposited at GEO under the accession number GSE207531. All other data is either included into the manuscript or available upon request.

