## Supplementary Figures 1-7 for "Constitutive production of flagellar proteins is required for proper flagellation in *Shewanella putrefaciens*"

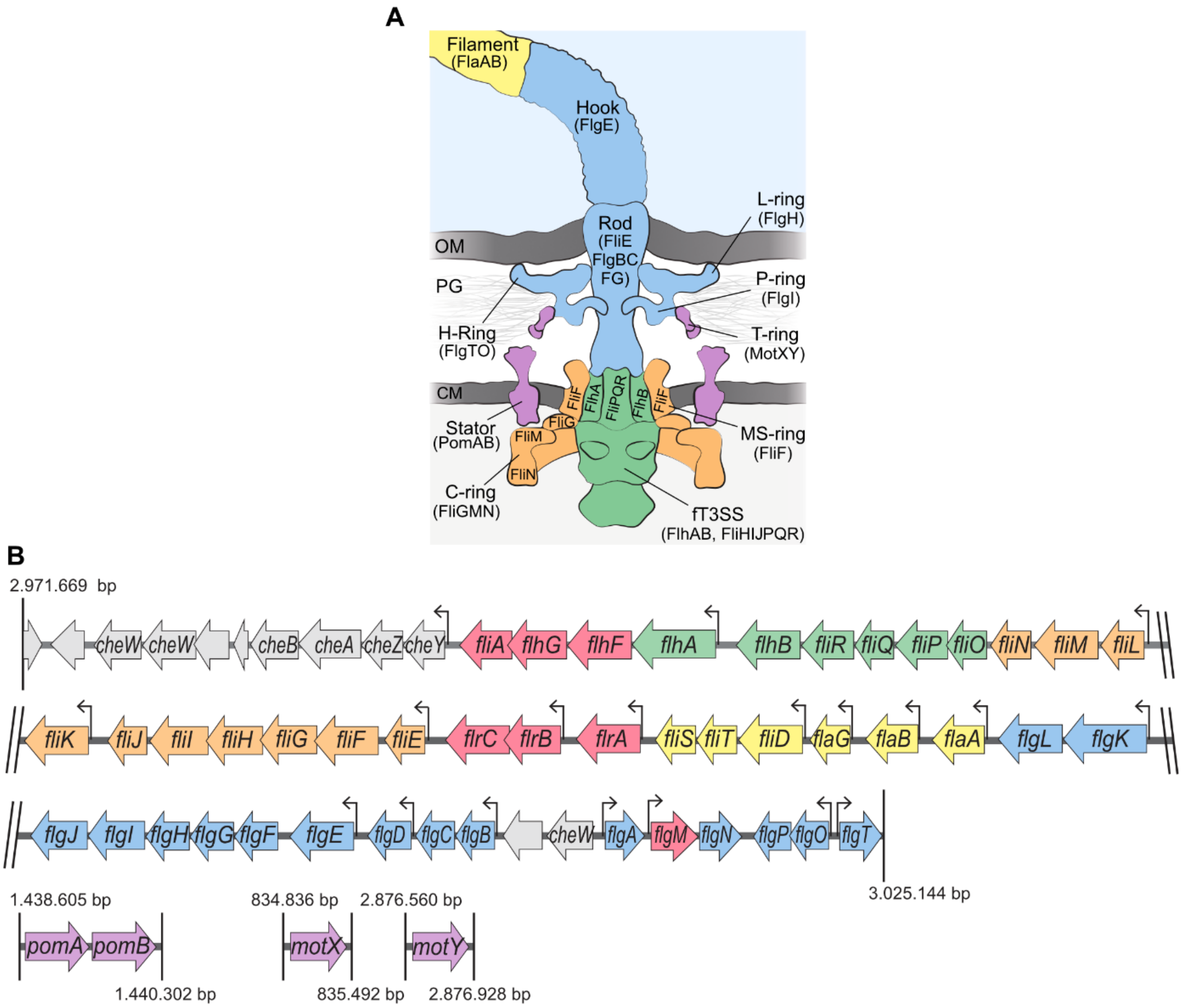


**Supplementary Figure 1: Composition of the *S. putrefaciens* polar flagellum and organizatin of the corresponding gene cluster. (A)** Schematic representation of a flagellum based on cryo-electron tomography images. The figure is modified from Kühn et al (2018) and Pecina et al (2021). **(B)** Genetic organization of the *S. putrefaciens* polar flagellar gene cluster of. Color coding of the genes corresponds to the colors of the components in the flagellar scheme. The promoters, as identified/predicted for *S. oneidensis* (Wu et al. 2011) are marked with arrows.

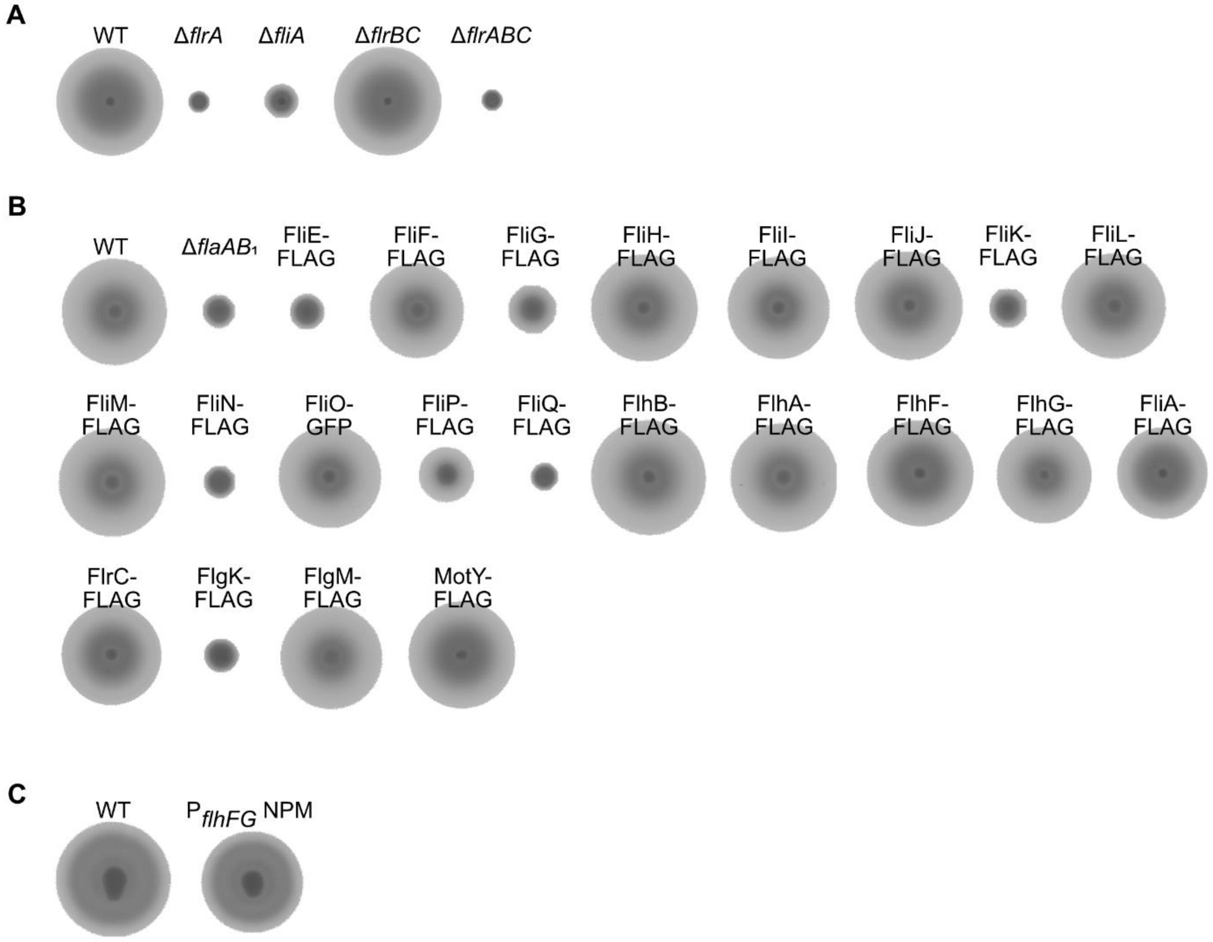


**Supplementary Figure 2: Spreading of *S. putrefaciens* strains in soft agar.** *S. putrefaciens* cultures with an $OD_{600}$ of approximately 0.5 were spotted onto LB soft-agar plates and incubated overnight at RT. The strains studied were always plated with the WT as a control on the same plate. **(A)** *S. putrefaciens* wild type spreading compared with that of *S. putrefaciens* deletion strains of the regulators as indicated. **(B)** Swimming abiltiy of the analysed *S. putrefaciens* strains with FLAG-tagged flagellar proteins. The deletion strain with polar flagellins (Δ*flaAB$_1$*) served as a negative control. **(C)** Swimming ability of *S. putrefaciens* bearing substitutions in the sequences of the $\sigma^{70}$-dependent promoters of *flhF* and *flhG* (NBS).

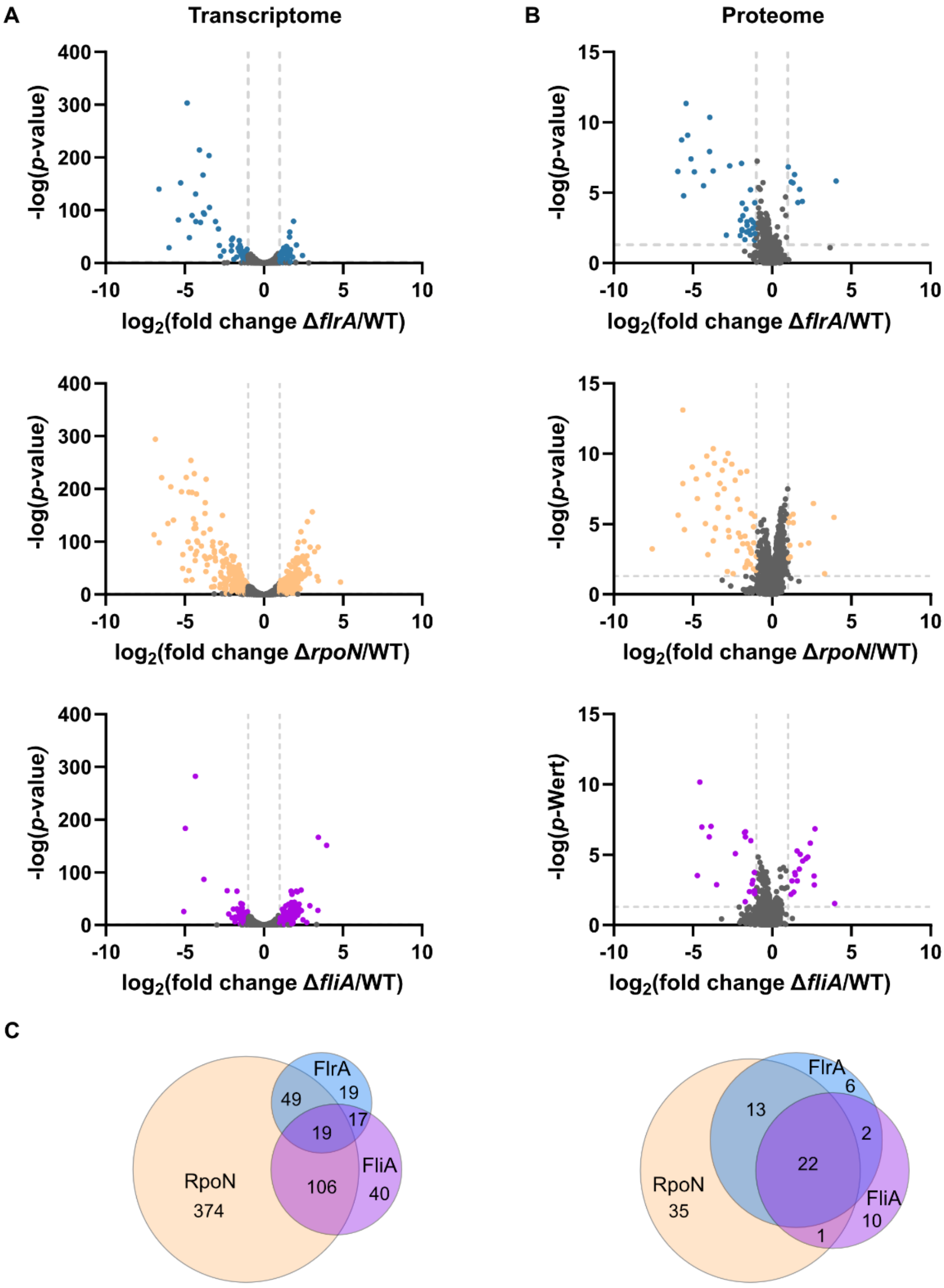


**Supplementary Figure 6: Affect on the transcriptome and proteome by FlrA, RpoN and FliA in *S. putrefaciens***: Volcano plots show the distribution of all genes **(A)** or proteins **(B)** with respect to their altered abundance ($\log_2$ fold change) and significance (-log *p*-value). The color-coded data points and dashed lines indicate the significantly changed mRNA and protein abundances, respectively. A $\log_2$-fold change of -1 and 1, respectively, and a p-value of 0.05 were assumed as significant. The altered transcriptome and proteome were analyzed in the Δ*flrA* (top, blue), Δ*rpoN* (middle, yellow), Δ*fliA* (bottom, purple) deletion strains compared with the *S. putrefaciens* wild type. **(C)** Absolute number of genes (left) and proteins (right) significantly altered in abundance comparing *S. putrefaciens* and deletion strains Δ*flrA* (FlrA), Δ*rpoN* (RpoN), and Δ*fliA* (FliA).

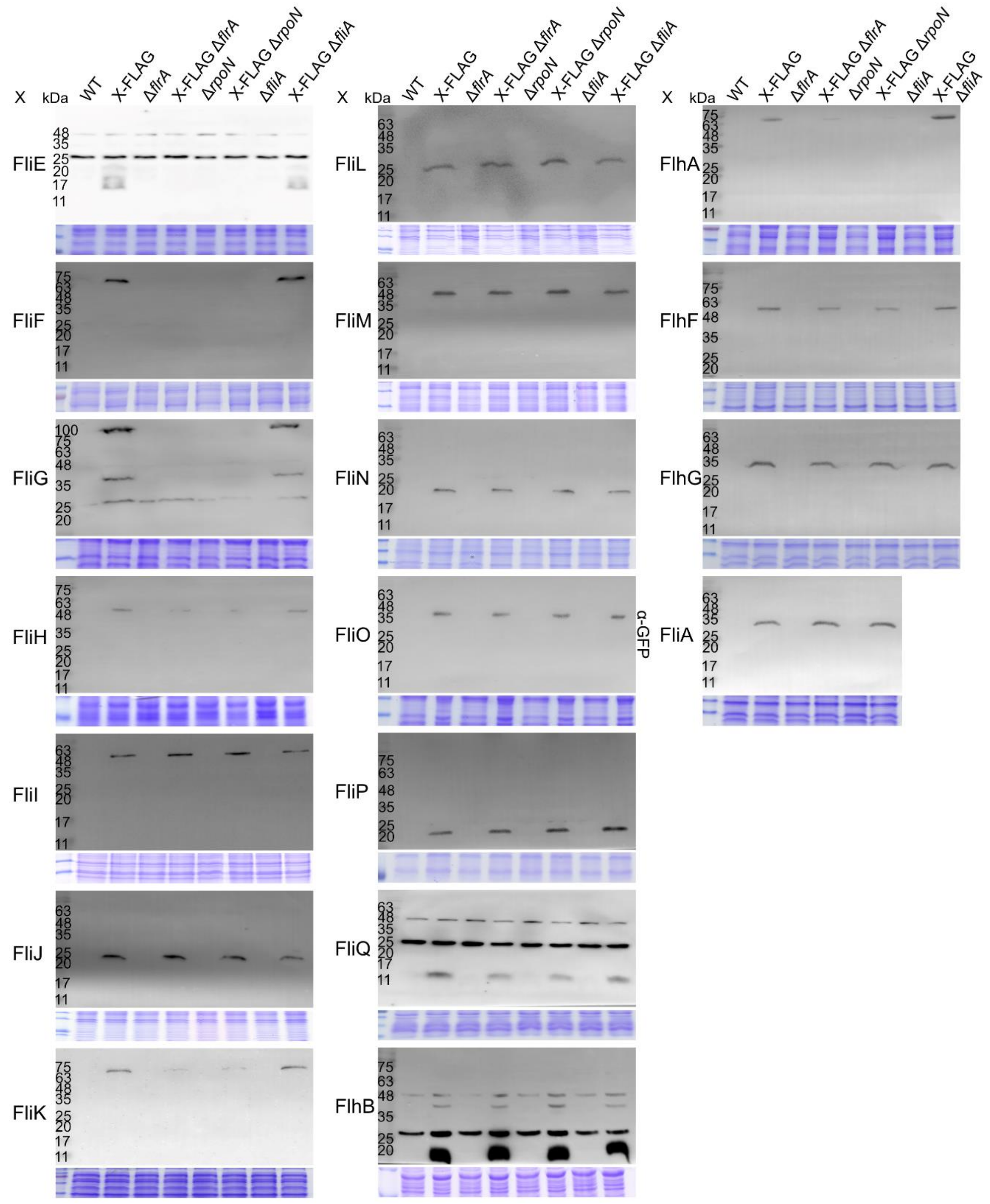


**Supplementary Figure 4: Western blot analyses on tagged flagellar proteins.** The figure displays the full blots shown in Figure 1 of the main manuscript. Most proteins were detected using an antibody against their FLAG fusion (α-FLAG-HRP). Detection of sfGFP-fused FliO was carried out using an antibody against GFP (α-GFP). The corresponding Coomassie-stained SDS-polyacrylamide gels served as loading control. Each experiment was conducted in biological triplicates.

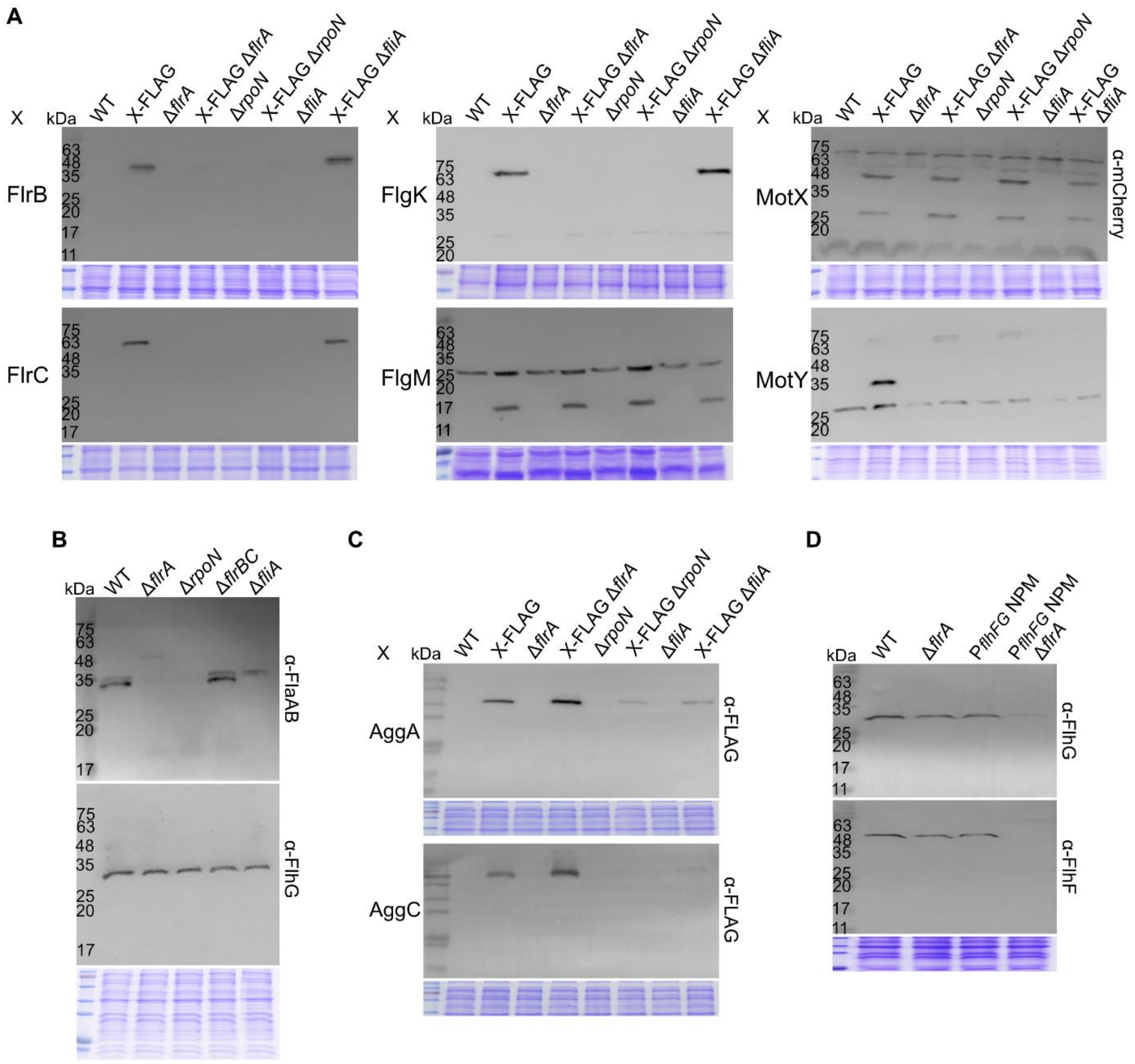

**Supplementary Figure 5: Western blot analyses on tagged flagellar proteins.** Display of the full blots shown in Figure 1 (**A** and **B**); Figure 3 (**D**) and Figure 4 (**C**) of the main manuscript. The proteins were detected using appropriate antibodies against a FLAG fusion (α-FLAG-HRP), an mCherry fusion (MotX; **A**, upper right panel; α-mCherry), the flagellins FlaA and FlaB (**B**), or FlhF and FlhG (**B** and **D**) . Each experiment was conducted in biological triplicates.

**A**

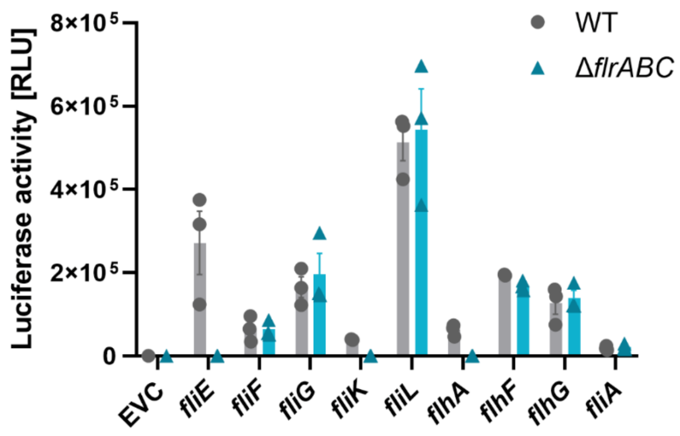


**B**

| **Gene** | **400 bp** | **Truncations** |
|---|---|---|
| EVC | 142 ± 7 | |
| *fliE* | 271680 ± 107216 | 86 bp: 77996 ± 13991 |
| *fliF* | *65023 ± 24892* | 100 bp: 404 ± 18<br>200 bp: 64495 ± 13223 |
| *fliG* | 165332 ± 35635 | 100 bp: 139065 ± 40048 |
| *fliH* | 149 ± 31 | |
| *fliI* | 295 ± 25 | 600 bp: 388 ± 22 |
| *fliK* | 40699 ± 1752 | 116 bp: 42851 ± 9812 |
| *fliL* | *513293 ±63059* | 100 bp: 930003 ± 4428<br>200 bp: 322208 ± 32921 |
| *fliQ* | 743 ± 183 | |
| *flhB* | 723 ± 19 | |
| *flhA* | 62320 ± 11496 | 80 bp: 104049 ± 55475 |
| *flhF* | 203671 ± 29840 | 100 bp: 147 ± 9<br>200 bp: 1448 ± 264<br>300 bp: 5496 ± 1851<br>550 bp: 194511 ± 1158<br>-284-550 bp: 205148 ± 7688 |
| *flhG* | 81872 ± 28137 | 100 bp: 634 ± 240<br>200 bp: 26717 ± 4976<br>300 bp: 24795 ± 2094<br>550 bp: 126136 ± 36668<br>-100-400 bp: 82349 ± 23616<br>-400-550 bp: 12621 ± 2060 |
| *fliA* | 18983 ± 4015 | 100 bp: 236 ± 88<br>200 bp: 327 ± 143<br>300 bp: 15604 ± 3177 |

**Supplementary Figure 6. Analysis of promoter activities using a luciferase reporter system: (A)** 400 or 550 bp fragments upstream of the translational start were fused to the reporter genes *luxCDABE* from *Photorhabdus luminescence*. These reporter plasmids with the promoter regions of interest were analyzed in the *S. putrefaciens* wild type and deletion mutant of the regulators *flr*ABC (Δ*flrABC*). **(B)** Truncated promoter versions were fused to the reporter genes *luxCDABE* from *Photorhabdus luminescence*. These reporter plasmids with the promoter regions of interest were analyzed in the *S. putrefaciens* wild type. Shown are the means and standard deviations from biological and technical triplicates.

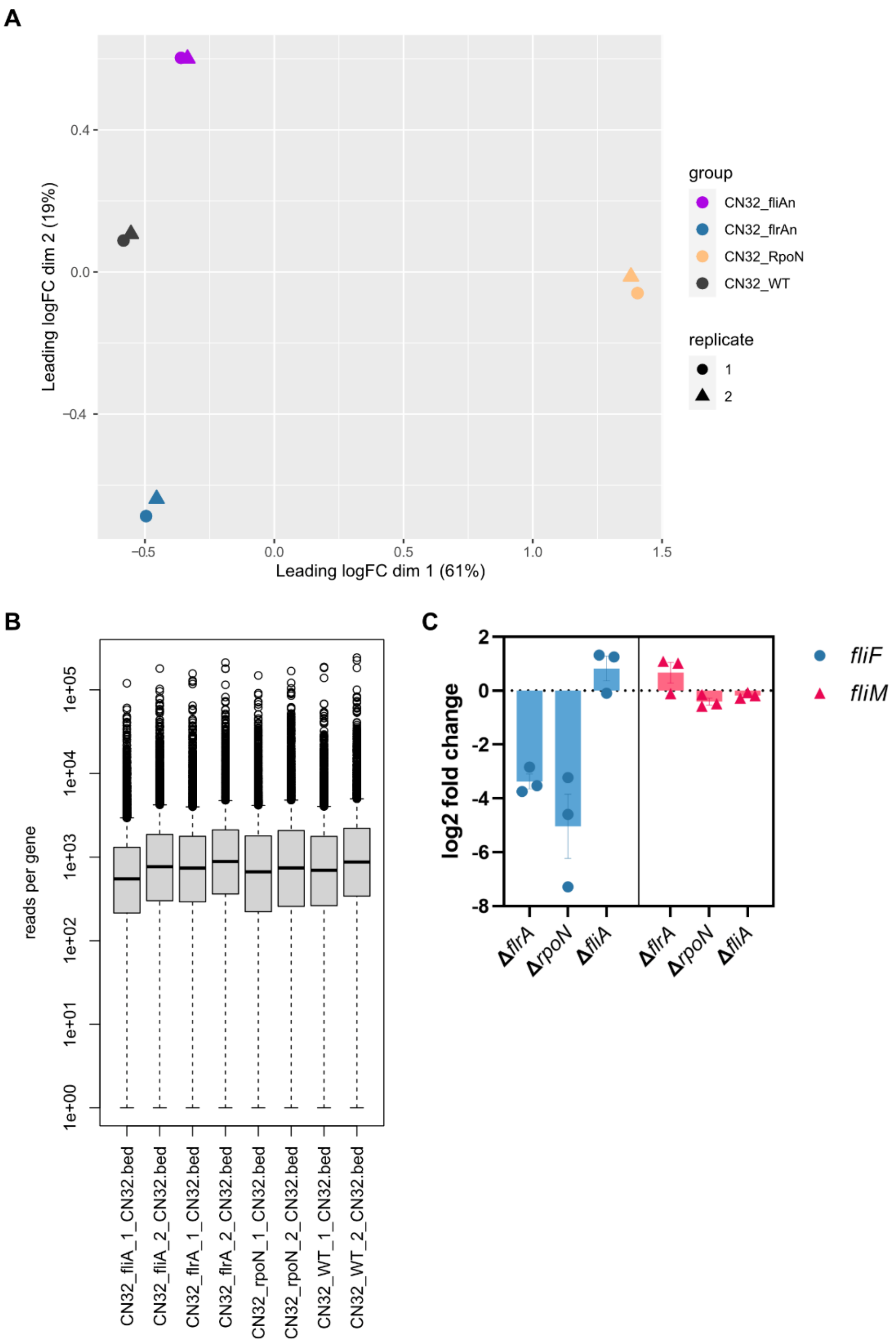


**Supplementary Figure 7: Transcriptome analysis reveals reproducible data generation for all analyzed strains. (A)** The *S. putrefaciens* wild type (CN32_WT) and the generated mutants (CN32_fliAn, CN32_flrAn, CN32_RpoN) were subjected to RNA sequencing. A multidimensional scaling analysis shows a clear separation of the transcriptional profile of the wild-type and the respective mutants, whereas the biological replicates for each strain cluster together. (**B**) A boxplot displaying the average reads per gene per strain reveal that across all samples the distribution of reads per gene are highly similar, which indicates that the growth conditions and handling of samples didn’t introduce a bias in the transcriptional profiles. **(C)** Differences of of *fliF* and *fliM* mRNA amounts in the indicated deletion strains could be reproduced by qRT-PCR. Means and standard deviations from technical duplicates and biological triplicates are shown.
